# DECODE: A *De*ep-learning Framework for *Co*n*de*nsing Enhancers and Refining Boundaries with Large-scale Functional Assays

**DOI:** 10.1101/2021.01.27.428477

**Authors:** Zhanlin Chen, Jing Zhang, Jason Liu, Yi Dai, Donghoon Lee, Martin Renqiang Min, Min Xu, Mark Gerstein

## Abstract

**Summary:** Mapping distal regulatory elements, such as enhancers, is the cornerstone for investigating genome evolution, understanding critical biological functions, and ultimately elucidating how genetic variations may influence diseases. Previous enhancer prediction methods have used either unsupervised approaches or supervised methods with limited training data. Moreover, past approaches have operationalized enhancer discovery as a binary classification problem without accurate enhancer boundary detection, producing low-resolution annotations with redundant regions and reducing the statistical power for downstream analyses (e.g., causal variant mapping and functional validations). Here, we addressed these challenges via a two-step model called DECODE. First, we employed direct enhancer activity readouts from novel functional characterization assays, such as STARR-seq, to train a deep neural network classifier for accurate cell-type-specific enhancer prediction. Second, to improve the annotation resolution (∼500 bp), we implemented a weakly-supervised object detection framework for enhancer localization with precise boundary detection (at 10 bp resolution) using gradient-weighted class activation mapping.

**Results:** Our DECODE binary classifier outperformed the *state-of-the-art* enhancer prediction methods by 24% in transgenic mouse validation. Further, DECODE object detection can condense enhancer annotations to only 12.6% of the original size, while still reporting higher conservation scores and genome-wide association study variant enrichments. Overall, DECODE improves the efficiency of regulatory element mapping with graphic processing units for deep-learning applications and is a powerful tool for enhancer prediction and boundary localization.

**Availability:** DEOCDE is available at decode.gersteinlab.org

## 1 Introduction

Transcriptional regulation in eukaryotes is the most common and fundamental form of gene regulation for maintaining cell identity during differentiation, determining how cells or organisms respond to intra- and extra-cellular signals, and coordinating various cellular activities (Cramer, 2019; Sperling, 2007). It undergoes precise spatial and temporal regulations via complex interactions of numerous cis-regulatory elements, such as enhancers and promoters, transcription factors (TFs), and chromatin remodelers (Abeel *et al*., 2009; Dao and Spicuglia, 2018; Jothi *et al*., 2009; Lewis *et al*., 2019; Klemm *et al*., 2019). Hence, enhancer discovery is the cornerstone for understanding transcription control and gene regulation.

The computational methods traditionally utilized for enhancer discovery mainly fall into two categories. First, some methods use the combinatory patterns of various epigenetic features within a genomic region (e.g., 200 bp bins) to infer the existence of enhancers with unsupervised approaches (Ernst and Kellis, 2012; Hoffman *et al*., 2012; Moore *et al*., 2020). These methods use unlabeled datasets to characterize chromatin states and require human interpretation of the discovered states. They are highly transferrable for predicting enhancers in a new cell type because the epigenetic patterns are dynamic and analogous across cell types. However, the simple combinatory, and usually binary, epigenetic patterns may not fully capture the complexity of transcriptional regulation. Second, some methods employ supervised approaches to identify target regulatory elements using hypothetical enhancer loci or a limited number of validated enhancers, which are underpowered for training a reliable model for accurate prediction (Alipanahi *et al*., 2015; Chen *et al*., 2018; Li *et al*., 2018; Lu *et al*., 2015; Min *et al*., 2017; Tang *et al*., 2020). We recently developed a linear predictive model based on shape-matching filters from multiple epigenetic features trained from genome-scale STARR-seq experiments on *Drosophila* (Sethi *et al*., 2020). However, most existing methods can only make binary predictions within a given region (>200 bp) and do not have a high enough resolution for more precise enhancer localization and boundary detection. Previous studies have shown that enhancers can range from 50-1,500 bp long (Dao *et al*., 2017; Li and Wunderlich, 2017). Hence, current enhancer annotations usually contain both active enhancer regions and redundant non-functional regions, introducing noise and reducing statistical power for downstream analyses such as casual variant mapping and functional validation.

Distinct from previous binary classification efforts, we reformulate the enhancer discovery problem into a weakly supervised **<u>object detection</u>** problem originated from computer vision by answering two questions: 1) Is there an enhancer within a given genomic region? 2) If yes, where is the enhancer? To accomplish these tasks, we take advantage of recent advances in functional characterization assays and utilize direct human enhancer activity readouts from STARR-seq experiments as fuzzy ground-truth labels (Muerdter *et al*., 2015). First, we propose a deep convolutional neural network (CNN) using epigenetic feature signals as input based on a simple but validated hypothesis – the magnitude, shape patterns, and cross feature coordination are important aspects of characterizing the identity of an enhancer (Schmidhuber, 2015). Concretely, we hypothesize that the interaction of open chromatin and histone marks provides a platform for TF binding, which allows epigenetic features to be predictive of enhancers (Mahony *et al*., 2005; Saeys *et al*., 2007; Spitz and Furlong, 2012). Second, we add a weakly supervised object detection module to precisely localize the target enhancer in the input genomic region. Specifically, we use the visual explanations created by gradient-weighted class activation mapping (Grad-CAM) for interpreting decisions from CNNs, which allows us to impute high-resolution enhancer coordinates that were never exposed to the model during training from fuzzy and coarsely labeled STARR-seq data (Selvaraju *et al*., 2017).

In the following sections, we describe our weakly supervised *<u>De</u>*ep-learning framework for *<u>C</u>*on*<u>de</u>*nsing enhancers and refining boundaries (DECODE) implemented in Python with TensorFlow. We performed extensive benchmarking using cell-line and transgenic mouse tissue validation data, and demonstrate that DECODE outperforms the *state-of-the-art* enhancer discovery models. We also validated the regulatory impact of our refined enhancer annotations using phylogenic conservation scoring, rare single-nucleotide polymorphism (SNP) enrichment, and genome-wide association study (GWAS) variant enrichment via stratified linkage disequilibrium score regression (LDSC).

## 2 Methods

We structured the task of cell-type-specific enhancer discovery as a weakly-supervised object detection problem with two modules. First, we constructed a **<u>CNN binary classifier to predict the existence of enhancers</u>**. The model takes a matrix of high-resolution epigenetic features over a large genomic window as input. Second, we developed an **<u>object detection module to locate the enhancer boundaries</u>** in the positive genomic windows based on the most informative subset of epigenetic features indicated by Grad-CAM. With a trained model, we can carry out cell-type-specific enhancer discovery in a novel cell type with common epigenetic profiles and obtain high-resolution core enhancer coordinates. We thoroughly benchmarked and validated our framework using various internal and external evaluation metrics. By evaluating through different biological perspectives, we demonstrate DECODE’s ability to generate high-quality cell-type-specific enhancer annotations with a strong regulatory impact.

### 2.1 Training data processing

We collected STARR-seq data for five human cell lines (HepG2, K562, A549, MCF-7, HCT116), along with chromatin accessibility (ATAC-seq and/or DNase-seq) and ChIP-seq for H3K27ac, H3K4me3, H3K4me1, and H3K9ac, from the ENCODE data portal (Appendix Table 1) (ENCODE Project Consortium and others, 2004). To call STARR-seq peaks, we applied STARRpeaker, which adjusts for GC content and RNA thermodynamic stability during peak calling (Lee *et al*., 2020). STARR-seq peaks overlapping with a chromatin peak and a peak of an active histone enhancer mark were defined as active enhancers and were considered positive training samples (Zhang *et al*., 2008). Negative regions were down-sampled from the background at a 1:10 positive to negative ratio. The positive and negative samples were extended to 4 kb, and the signals were aggregated over 10 bp bins.

**Table 1.** DECODE training and testing performance: Out-of-sample validation performance metrics for each chromatin accessibility and cell type.

| Chromatin Accessibility | Cell Type | Validation Accuracy | Validation auROC | Validation auPRC |
| --- | --- | --- | --- | --- |
| ATAC-seq | K562 | 0.9885 | 0.9972 | 0.9704 |
|  | HepG2 | 0.9908 | 0.9960 | 0.9536 |
| DNase-seq | K562 | 0.9849 | 0.9984 | 0.9975 |
|  | HepG2 | 0.9865 | 0.9978 | 0.9972 |
|  | A549 | 0.9818 | 0.9984 | 0.9978 |
|  | HCT116 | 0.9918 | 0.9989 | 0.9981 |
|  | MCF-7 | 0.9897 | 0.9983 | 0.9978 |

The resolution of signal aggregation determines the precision of boundary detection. Here, every value in the input matrix represents the average epigenetic signal of a 10 bp bin. In the end, each input value is assigned a Grad-CAM importance score. Hence, filtering by the importance score obtains core enhancers at a 10 bp resolution. It is possible to extract higher-resolution signals for higher precision, but 10 bp was the experimentally determined optimum resolution and the highest resolution for most ChIP-seq experiments.

For the ATAC-seq version, there were 211,097 STARR-seq peaks and 459,321 ATAC-seq peaks for both (HepG2, K562) cell types. Only 25,420 of the STARR-seq and ATAC-seq overlap regions intersected with another ChIP-seq, which were selected as the positives. For the DNase-seq version, there were 912,967 DNase-seq peaks for all five cell types. Only 73,271 of the STARR-seq and ATAC-seq overlap regions intersected with another ChIP-seq, which were selected as the positives. For the selected positive regions, we observed a distinct signal shape for each assay. The peak in chromatin accessibility and peak-trough-peak in other histone marks validate our selection for the training regions and provide a basis for CNN pattern recognition (Fig. 4).

### 2.2 Binary classifier construction

As shown in Fig. 1, the model is a ResNet-inspired CNN that contains convolutional layers, pooling layers, and dense fully connected layers (He *et al*., 2016). The input is a data matrix (of size 5×400) containing values from the signal tracks of the five epigenetic assays extracted from a 4 kb region by aggregating the signals over 10 bp bins. Each value in the input matrix represents the signal of an epigenetic assay at a genomic location. We use only epigenetic features because they are more generalizable compared to sequence-based features, especially when transferring predictions to unseen cell types (Zhou *et al*., 2011). Our model contains seven convolutional layers, each of which uses its *k* convolutional filters to produce *k* activation maps of width *i* and height *j*: **A**^*k*^ ∈ ℝ_*i*×*j*_ with weights **W**^*k*^ and bias **B**^*k*^ from an input **X** in layer *l*.

**Figure 1.**
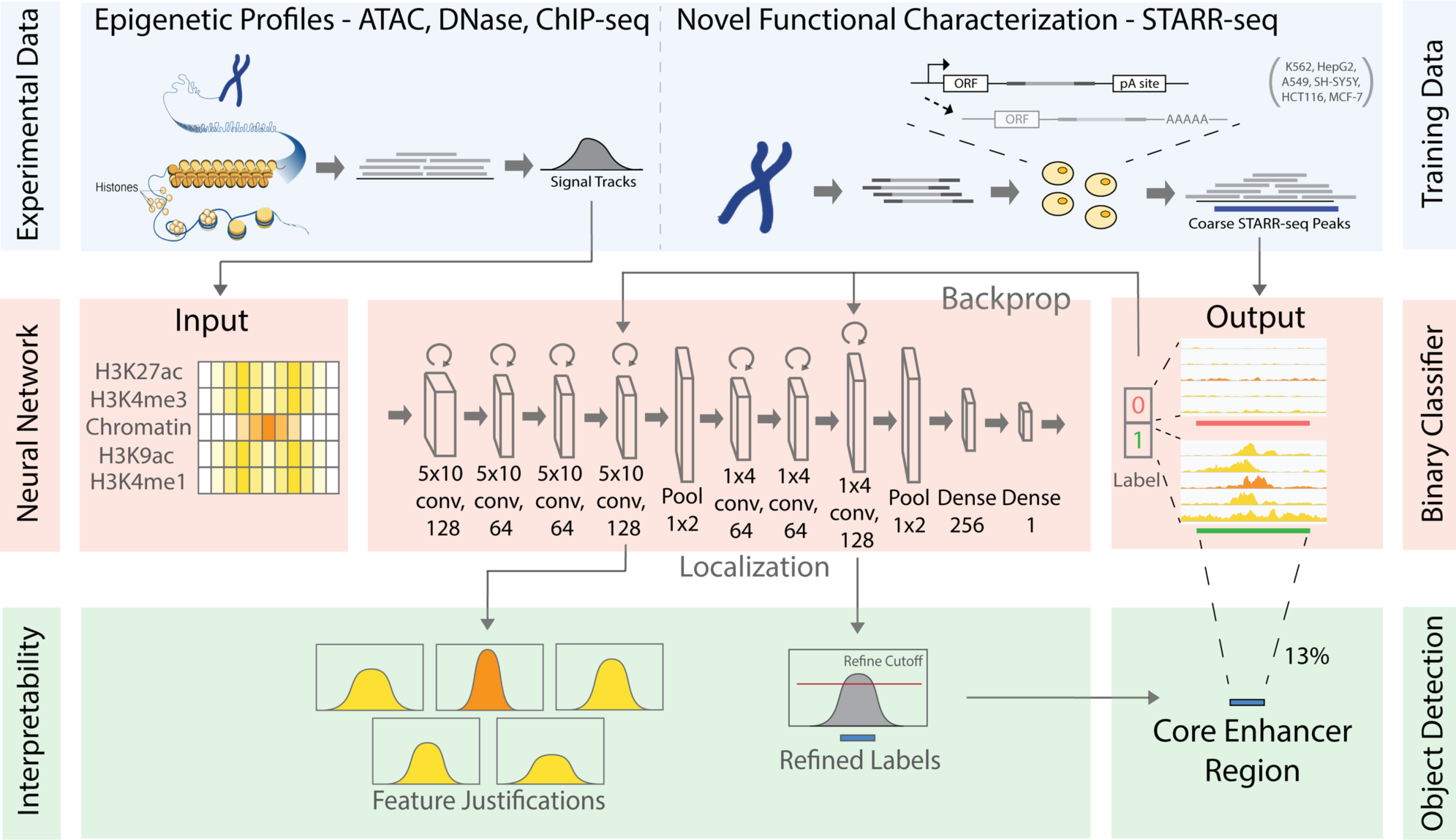
DECODE Model Schematics: DECODE has three major components. First, the model uses epigenetic features and low-resolution STARR-seq peaks as training data. Second, it uses the epigenetic profiles to predict the presence of enhancers with a CNN. The architecture is composed of two sets of convolution-pooling layers followed by two dense layers. The input consists of a matrix of signals from five epigenetic experiments. The output is a sigmoid probability of the 4 kb input region containing enhancers. Third, feature-wise and position-wise Grad-CAM scores are calculated for interpretability and boundary detection. Position-wise Grad-CAM scores are used to extract the core enhancer regions.

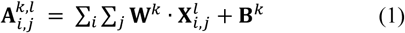

The first several convolutional layers extensively capture altitude and shape-based features from either chromatin accessibility or ChIP-seq with different convolution filters. Moreover, the filters blend signals across different tracks to allow for combinatorial feature extraction. Then, max-pooling layers are used to reduce the number of parameters and abstract features trained in the previous convolutional layers. If *h, w* denotes the dimensions of a pooling operation, then max-pooling over the activation map takes the max value of each *h, w* window to produce an output of size *m* = *i*/*h* and *n* = *j*/*h* as input for the next layer.

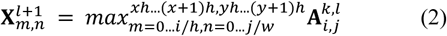

We hypothesize that the interaction of open chromatin states and histone marks coordinate TF binding in enhancers (Spitz and Furlong, 2012). Therefore, we placed chromatin accessibility in the middle of the input data matrix to allow for maximal interaction with other histone marks. The first few filters have a kernel size of 5×10 in order to span all five assays. With padding removed, the features are then convoluted and down-sampled to one dimension to represent linear genomic windows. Further convolutions on the linear feature use a one-dimensional (1D) filter of size 1×4. The pooled layers are then fed into fully connected layers to make a sigmoid prediction on the probability of enhancers being in the region.

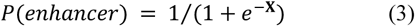

Between every convolution layer are squeeze-and-excitation blocks that calculate the residual features and act as a gate for how much original feature is passed through (Hu *et al*., 2018).

### 2.3 Object boundary detection via weakly supervised learning framework (Grad-CAM)

We operationalized the task of enhancer localization using a weakly supervised object detection method in computer vision. Grad-CAM extracts the implicit localization of the target from classification models and obtains a high-resolution subset of the image with the most informative content regarding the target (Selvaraju *et al*., 2017). For a genomic region with a positive classification, we used Grad-CAM to extract the implicit enhancer localization as a subset of the original input genomic region, thereby increasing the resolution of our core enhancer annotations (Fig. 2). Utilizing Grad-CAM to revisit the positive predictions can refine our annotations by finding the most salient enhancer regions. Further, our method is much more interpretable compared to previous supervised black-box prediction models because we can trace and visualize the process of decision-making in our network.

**Figure 2.**
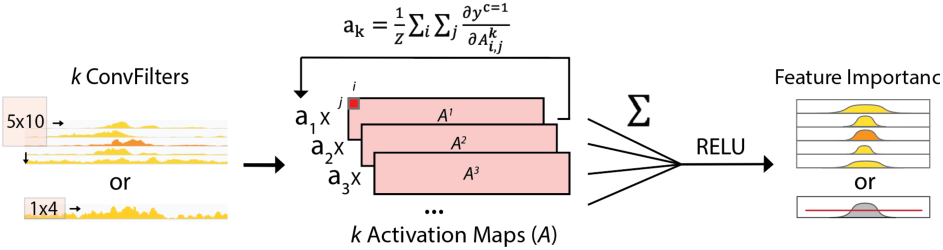
Grad-CAM for Enhancer Localization: After a positive prediction, we can use Grad-CAM to extract epigenetic features and genomic locations that were important in making the positive prediction. By superimposing activation maps (*A*^*k*^) weighted by an importance score (a_k_), Grad-CAM maps highlight the most salient features in making a final output, which we used to generate high resolution core enhancer annotations.

During the training process, each convolutional filter learns to extract features that are important in accurately predicting enhancers in the genomic window and outputs an activation feature map. Each activation feature map highlights genomic regions that contain important features for enhancer prediction. Superimposing all activation feature maps sums together all the highlighted regions and forms a silhouette of locations, with respect to the original input, that activated the greatest number of neurons in our neural network, thereby extracting the implicit localization information underlying a classification task.

In detail, we produce a scalar importance score for each activation map using the global-average-pooled gradient of the positive class with respect to the feature map activation (Equation 4).

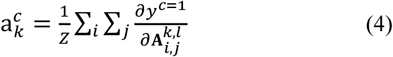

We use the scalar importance scores as weights for the linear combination of all *k* activation feature maps. We then multiply each importance score with its respective activation feature map and sum element-wise over all activation feature maps in the layer. Next, we apply RELU to filter for only positive values, and interpolate to obtain a Grad-CAM score map of size *i* × *j*, the same size as the input of the convolutional layers (Equation 5).

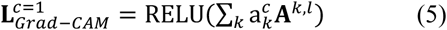

Grad-CAM maps can be generated for different convolutional layers in our architecture. Concisely, the Grad-CAM map from the last 1D convolutional layer is a 1D map, or a position-wise score, describing the highest-level feature importance of each 10 bp bin in the genomic window. To refine our predictions, we use the position-wise score to select a subset of the input region (or a subset of the 10 bp bins) that corresponds to a high Grad-CAM score. Here, our cutoff is set as the average Grad-CAM score over all positive genomic positions; hence, the filtering is performed after all predictions have been processed by Grad-CAM. Furthermore, Grad-CAM maps can also be generated using gradients from the first few convolutional layers to observe how much each epigenetic assay contributes to a positive prediction, or a feature-wise score. We constructed our custom CNN such that the high- and low-level Grad-CAM maps correspond to position-wise or feature-wise biological interpretations, respectively. This enables us to study the epigenetic features of enhancers by visualizing and interpreting the process of decision-making of our model.

### 2.4 Model training configurations

We used the Adam optimizer at a learning rate of 5e-5 and added dropout layers to prevent overfitting. The model is set to train for 100 epochs, but we included early stopping with monitoring of validation loss to prevent overfitting. The data was split 80-20 for training and validation. We also added class weights to address the 1:10 positive and negative sample imbalance in our dataset. Training and model prediction were accelerated using NVIDIA Tesla K80 GPUs.

### 2.5 Enhancer challenge data for benchmarking against the state-of-the-art model

We used transgenic mouse enhancer data from the VISTA enhancer database to benchmark against the state-of-the-art method (Visel *et al*., 2007). Hypothesized enhancers were cloned into a plasmid with a promoter and *lacZ* reporter gene, which were injected into mouse embryos. After reimplantation with surrogate mothers, the transgenic embryos at e11.5 to score for enhancer activity. Target regions were considered positive for enhancers if at least three transgenic embryos had reporter-gene expression across three sample tissues and were considered negative if we observed no reproducible pattern across at least five samples. Tissue-specific enhancers were pooled from all six tissues (forebrain, heart, hindbrain, limb, midbrain, neural tube). In order to mediate cross-species effects, DECODE was trained with all available human cell line data, and then fine-tuned with out-of-sample mouse enhancers.

### 2.6 Cell-line case study evaluation

We utilized a variety of methods to evaluate the accuracy of our supervised model and the regulatory impact of our core enhancer annotations. As a case study, we predicted enhancers in neural progenitor cells (NPCs), which lack STARR-seq data. These cells play an important role in psychiatric disorders.

The NPC signal tracks for DNase-seq and four active histone marks (H3K27ac, H3K4me3, H3K9ac, H3K4me1) were downloaded from the ENCODE portal. We created sliding windows of size 4 kb with 500 bp steps across the whole genome. For DECODE classifier input, the average signal was extracted for 10 bp bins in each genomic window. After classification, the positive 4 kb genomic windows made up the original annotation. We then applied Grad-CAM to refine each positive 4 kb window to define our core enhancer annotation.

#### GWAS LDSC Enrichment

We characterized the disease-variant impact of our NPC annotations by calculating stratified LDSC enrichment from GWAS (Bulik-Sullivan *et al*., 2015). LDSC regresses the chi-square statistics (*X*^2^) with the linkage disequilibrium (LD, *r*^2^) to estimate the heritability in a disease-specific manner. This method calculates the partitioned heritability of certain regions or annotations using GWAS summary statistics. We utilized LD scores from the 1,000 Genomes Project and GWAS summary statistics from Bulik-Sullivan *et al*. and the Psychiatric Genomic Consortium (Siva, 2008; Turley *et al*., 2018).

#### Conservation Score Analysis

We measured inter- and intra-specie conservation by 100-way PhastCons and rare derived allele frequency (DAF) SNP enrichment, respectively. The PhastCons score is a phylogenetic hidden Markov model trained on genetic sequences across 100 different species and quantifies the conservation of a given genetic annotation (Yang, 1995). We calculated the ratio of DAF (<0.5%) SNP enrichment using SNPs from the Genome Aggregation Database (gnomAD) and Pan-Cancer Analysis of Whole Genomes (PCAWG) resources (Campbell *et al*., 2020; Karczewski *et al*., 2020).

## 3 Results

In contrast to traditional enhancer classification methods, we reformulated enhancer discovery into a two-step weakly supervised object detection problem. Specifically, we first utilized the direct enhancer activity readout from novel functional characterization assays to train a deep neural network. Result sections 3.1 and 3.2 demonstrate the ability of our deep learning binary classifier to make accurate predictions. Then, for the object detection module, we used Grad-CAM to define high-resolution enhancer boundaries; sections 3.3 to 3.6 describe our results for enhancer localization. In short, we applied our two-step DECODE model to various real-world datasets for comprehensive performance benchmarking and demonstrate its benefits in constructing compact genome annotations to facilitate variant interpretations.

### 3.1 The DECODE binary classifier is a transferrable model for accurate cell-type-specific enhancer prediction

A trained binary classifier can predict enhancers on sliding windows across the genome. We merged and shuffled positive- and negative-labeled data from all cell types to train a binary classifier for predicting the existence of enhancers in any given 4 kb genomic region using cell-type-matched epigenetic features (details in Methods 2.4). To verify our results, we performed five-fold cross-validation by partitioning the merged data into five folds and iteratively using each fold as the out-of-sample validation set. High out-of-sample area under the receiver operating characteristic curve (auROC; ATAC-seq: 0.999, DNase-seq: 0.998) and area under the precision-recall curve (auPRC; ATAC-seq: 0.972, DNase-seq: 0.989) metrics demonstrate that the binary classifier module in DECODE can accurately predict enhancers using combinatory epigenetic features. In addition, we did not observe any divergence between training and validation loss during backpropagation. Moreover, the validation metrics remained high across all folds (>0.95), suggesting that there was no overfitting under our training configurations.

In real-world scenarios, a model would predict enhancers in cell types that it has not yet seen during training. Therefore, we further tested the robustness of the DECODE binary classifier in transferring the learned features to new cell types. In other words, we evaluated whether the high validation metrics are specific only to the cell types used in training or whether they can be easily generalized to other cell types. Hence, we performed cross cell-line validation by leaving out one cell type for validation while training on the rest of the cell types (for both ATAC-seq and DNA-seq). Similar to the validation performance trained from merged cell type data, we observed consistently high cross-cell-type validation metrics in all cell types, demonstrating our model’s ability to transfer its predictions across cell types (Table 1). For instance, our DECODE model trained on DNase-seq and four other ChIP-seq datasets achieved consistently high validation auROC (0.996-0.999) and auPRC (0.954-0.998) scores. The performance remained high even when using ATAC-seq for chromatin accessibility. Therefore, our DECODE model can accurately predict enhancers and robustly transfer the predictions onto a novel, unexplored cell line.

### 3.2 DECODE outperforms the existing *state-of-the-art* method on experimentally validated mouse enhancers

In addition to making accurate predictions for internal evaluations, we further compared the efficacy of our model with existing methods on an external experimentally validated dataset. To do so, we applied DECODE binary classifier on 3,244 experimentally validated regions from six mouse tissues, which were also used in the official ENCODE enhancer challenge. Specifically, we downloaded the signal tracks for the five epigenetic features from the ENCODE portal, extracted the signals for each given region as input, and predicted for the existence of enhancers using our trained model. We compared our predictions with Matched-Filter, the leading method in the ENCODE enhancer challenge (Sethi et al., 2020).

As shown in Fig. 3, our DECODE model obtained an average auPRC of 0.46, which was 24% higher than the auPRC of Matched-Filter on the same dataset (Fig. 3a). It is also worth pointing out that DECODE outperformed Matched-Filter in all six tissues with a decent margin. Specifically, our model demonstrated higher auROC scores ranging from 0.82-0.85 in all six tissues (vs. 0.76-0.85 for Matched-Filter) and noticeably improved auPRC scores (0.39-0.57 in DECODE vs. 0.27-0.43 in Matched-Filter) with an average margin ranging from 0.02-0.18.

**Figure 3.**
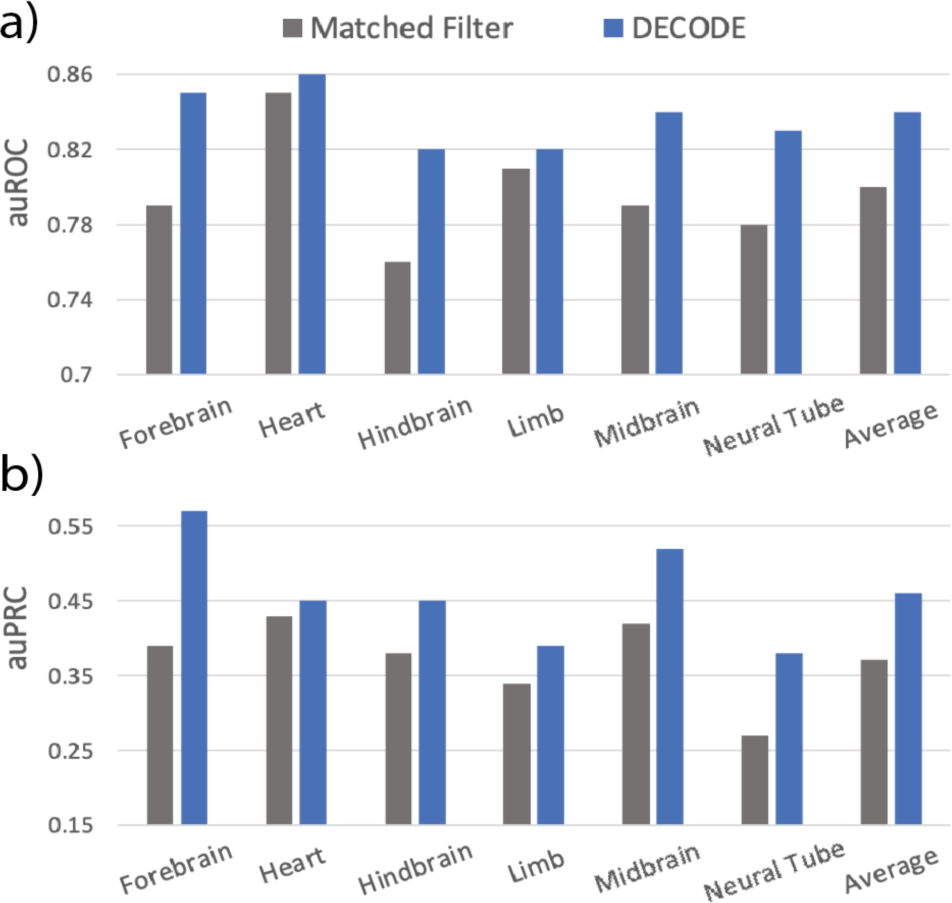
Benchmarking Against Matched Filter-Based Model: We benchmarked our trained model against the state-of-the-art model using transgenic mouse enhancers. DECODE produced validation metrics (a, auROC; b, auPRC) that outperformed the state-of-the-art model in all tissue types.

**Figure 4.**
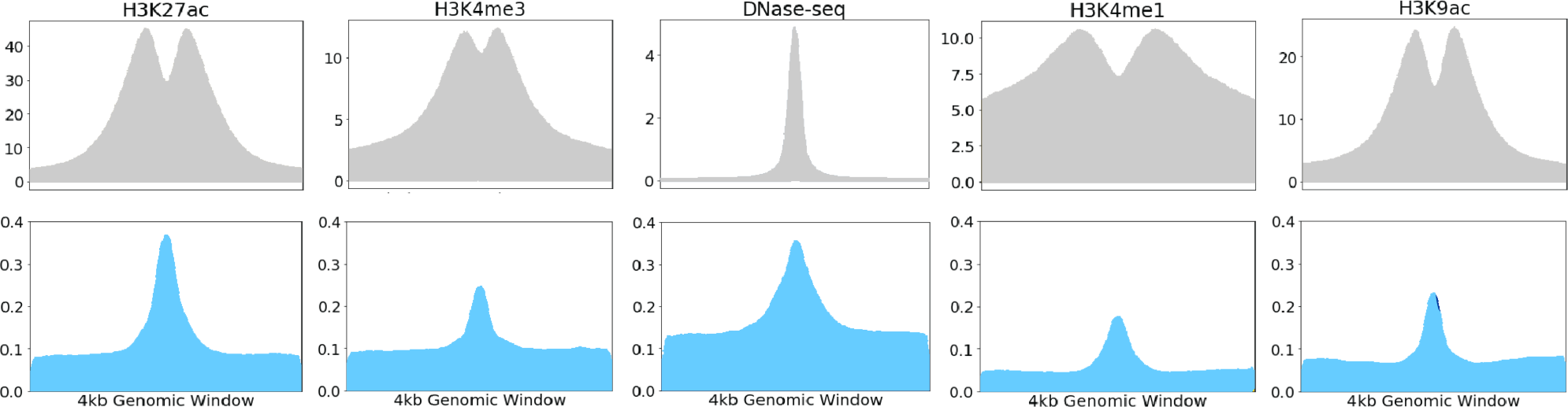
Feature-wise Grad-CAM score: Original signal (top row) and feature-wise Grad-CAM score (bottom row) over a 4 kb window for the five types of input epigenetic marks.

Two main reasons could explain DECODE’s improvement in performance over existing methods. First, we used genome-wide large-scale training data from direct human enhancer readouts of five cell types compared to the *Drosophila* data used for the training of Matched-Filter. This allows our model to recognize more complicated features that are important for enhancer predictions. Second, we believe the interactions among epigenetic features guarantee active regulatory activity in functional enhancer regions, which is demonstrated in previous literature (Mahony *et al*., 2005; Saeys *et al*., 2007; Spitz and Furlong, 2012). As a result, we designed the convolutional filters in our deep learning framework to span multiple epigenetic marks to model non-linear epigenetic interactions. In contrast, Matched-Filter considers different epigenetic marks independently with its linear support vector machine-based methods. The capacity of DECODE to learn complex combinations of features provides the basis for achieving better performance than the current state-of-the-art method.

### 3.3 DECODE’s object detection module generates interpretable visual explanations for enhancer boundary refinements

After demonstrating the efficacy of DECODE’s binary classifier for accurate enhancer predictions, we seek to uncover more information regarding the basis of neural network decisions through our weakly supervised framework. The DECODE object detection module extracts interpretable feature-wise and position-wise importance scores as a visual explanation. Here, we examine the benefits of feature-wise importance scores generated by Grad-CAM from the lower-level convolutional layers.

Grad-CAM uses gradients to identify input locations that activate the greatest number of neurons in a given layer. We extracted the feature-wise Grad-CAM scores from the positive training samples to assess the basis of model predictions. A high feature-wise Grad-CAM score corresponds to higher importance placed on that epigenetic feature. Grad-CAM scores peak at the center of the 4 kb windows for each epigenetic assay (Fig. 4), which corresponds to the greatest amount of regulatory activity as indicated by the original signal. This result shows that DECODE predicts the presence of enhancers using highly active regulatory regions.

Next, we compared the Grad-CAM scores across different epigenetic features to show that our model prioritizes key features for enhancer prediction. As shown in Fig. 4, DNase-seq and H3K27ac ChIP-seq demonstrate the highest Grad-CAM scores (mean at center >0.35) as compared to other features (mean at center <0.25), indicating their important role in defining enhancers in the genome. Our finding recapitulates known biology, as these features were also used in the official ENCODE3 encyclopedia annotation (Moore *et al*., 2020). In contrast, H3K4me1 exhibited the lowest feature-wise Grad-CAM score (mean from 0.05-0.2), which implies that it played a relatively less important role in our model decision.

### 3.4 DECODE provides high-resolution enhancer boundary localization

Position-wise Grad-CAM importance scores from DECODE’s object detection module can be used to interpolate high-resolution cell-type-specific enhancer coordinates and condense the annotation of an enhancer to its core functional regions. In contrast, this feature is missing in most existing methods.

To demonstrate this function, we utilized DECODE to predict compact enhancers in NPCs, which play important roles in neurodevelopment and have been implicated in a wide variety of psychiatric disorders (Castrén, 2014; Das *et al*., 2020). Specifically, we utilized the five epigenetic marks on NPCs to predict enhancers using our DECODE framework trained on all available data (see details in Methods 2.4). We divided the genome into 4 kb windows with 500 bp steps between each window. The corresponding epigenetic signals were extracted at a 10 bp resolution from each window as inputs to our model. Windows with a binary classifier output greater than 0.5 were identified as positive predictions. In total, we identified 1,515,431 overlapping windows across the genome, and 1.1% (17,622) among them showed predicted enhancer activities. We merged the positive predictions to create the original prediction set, which contained 16,522 elements with a mean length of 4,188 bp.

To condense the annotation using DECODE’s object detection module, we extracted the position-wise Grad-CAM scores from the positive regions for each 10 bp bin (Fig. 6a). Bins with Grad-CAM scores larger than the mean across all windows were merged to create the refined set. This process resulted in a total of 23,505 refined positive elements with a mean length of 371 bp.

In Fig. 5, we show an example positive genomic window (chr1:8680000-8684000) and its feature-wise and position-wise Grad-CAM values. The example shows the process by which high-level position-wise scores are derived from low-level feature-wise scores, which increases interpretability by tracking neuron activation through our neural network classifier.

**Figure 5.**
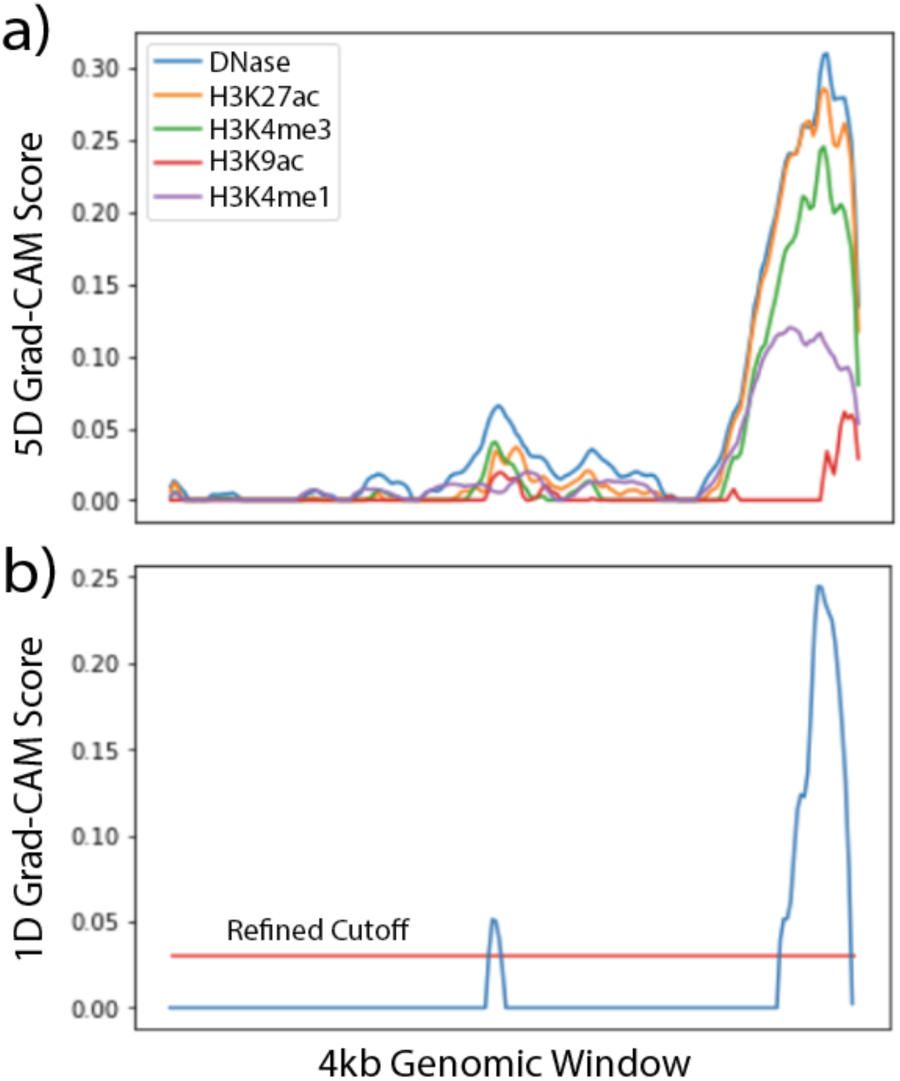
Feature-wise and Position-wise Grad-CAM Values: a) 5-D and b) 1-D Grad-CAM justification of a positive prediction.

**Figure 6.**
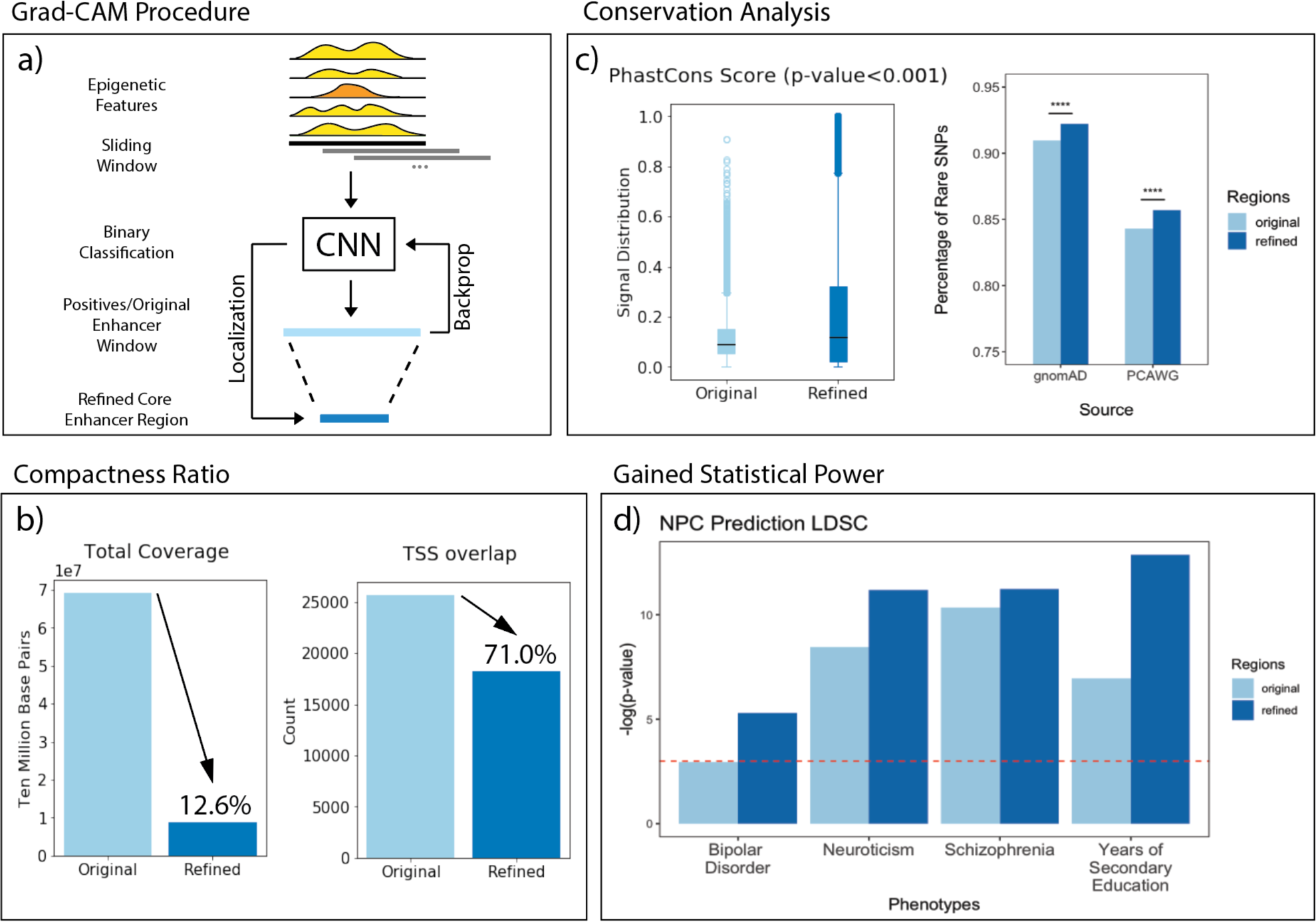
NPC Whole Genome Prediction Validation: a) Procedure to predict enhancer windows and generate refined regions. b) Total nucleotide coverage and total transcriptional start site overlap for the original versus refined set. c) Conservation analysis of the PhastCons score distribution and the rare DAF SNP enrichment of the refined set compared with original set. d) LDSC enrichment of psychiatric and neurodevelopmental phenotypes for the original and refined set.

In addition, the true enhancer in the example is shifted to the right of the genomic window. Due to the lack of a quantitative boundary detection algorithm, most existing methods take the entire input window as an enhancer region, which potentially confounds various downstream analyses such as validation region selection and disease causal variant mapping. In contrast, the object detection module in our DECODE model does not rely on interpolating from the center, but rather localizes enhancer coordinates based on the importance of the loci using the Grad-CAM outputs. Therefore, we are still able to rescue the shifted enhancers and discover true functional regions with high regulatory impacts.

This boundary detection module can remove a significant portion of the noise regions in our enhancer prediction and noticeably condense our genome annotation. For instance, analysis on the refined set shows that it is only 12.6% in coverage as compared to the original set but includes a disproportionately large amount (71%) of transcription start sites, which indicate a more enriched transcriptional regulatory footprint (Fig. 6b). In the following sections, we demonstrate the compactness of the refined set through conservation and GWAS variant enrichment analysis.

### 3.5 DECODE’s compact enhancer predictions are highly conserved across species and populations

To test whether the object detection module accurately selects true regulatory regions, we compared cross-species conservation scores of the original positive input regions vs. condensed core enhancer regions. If the enhancer annotations serve important regulatory functions, then those annotations should be conserved, as any negative mutations would increase the likelihood of disadvantage phenotypes that eliminate that allele from the gene pool. Hence, higher conservation usually indicates higher enhancer quality (Yang, 1995).

Specifically, we downloaded the 100-way PhastCons scores and calculated the average PhastCons conservation scores within each original and refined region. The median PhastCons score in the refined core regions was 0.117, which is significantly higher than those in the original positive regions (median 0.092, one-sided Wilcoxon test P < 0.001, Fig. 6c).

We further evaluated the quality of our compact enhancer annotation from DECODE via the enrichment of rare variants with a simple but validated assumption – key functional regions in the genome are under strong negative selection and hence are depleted in common variants (Fu *et al*., 2014; Zhang, Liu, *et al*., 2020). Therefore, we compared the rare variant enrichment in the refined and original positive enhancer regions. Specifically, we downloaded the entire human genetic variation set from gnomAD and PCAWG (Campbell *et al*., 2020; Karczewski *et al*., 2020), and defined rare variants as those with a DAF less than 0.5% over the entire population. We calculated the percentage of rare variants within each merged annotation set. The refined enhancer regions demonstrated significantly higher percentages of rare variants in both the gnomAD and PCAWG dataset. For instance, we observed a rare variant percentage of 0.922 and 0.857 in gnomAD and PCAWG, respectively; this number decreased to 0.910 and 0.843 for the original positive input regions (P < 10^5877.9^ for binomial test in both datasets).

Altogether, the condensed compact enhancers refined in our model showed higher cross-species and cross-population conservations, indicating DECODE’s ability to remove noisy regions in our enhancer predictions and provide high-quality genome annotations.

### 3.6 Compact enhancer annotations predicted by DECODE can better explore GWAS variants in psychiatric disorders

Disease-causal variant mapping is one of the most important applications of distal regulatory element mapping. Lines of evidence have demonstrated that accurate and compact annotations can significantly increase the statistical power for both somatic and germline variant mapping in disease studies (Fu *et al*., 2014; Zhang, Liu, *et al*., 2020; Zhang, Lee, *et al*., 2020). Therefore, we further tested whether our condensed enhancer definitions can benefit variant prioritization and interpretations.

Here, we predicted two set of enhancers in NPCs – a coarse set of predictions using the binary classifier (similar to existing methods) and a refined set of core predictions using the DECODE object detection module. We extracted the summary statistics from GWAS for around 2 million SNPs for four NPC relevant traits – bipolar disorder, neuroticism, schizophrenia, and years of secondary education. For each phenotype, we used stratified LDSC to test whether the heritability of a phenotype is enriched in one set of annotated genome regions in NPCs, where a high LDSC enrichment for a GWAS trait would indicate that the set of annotations has a high partitioned heritability for the corresponding trait. Using these summary statistics, we calculated the enrichment of the original and refined set for each trait, represented by the P-value (Fig. 6d). We found that both datasets demonstrated significant LDSC enrichment for three of the four phenotypes (with log P-value ranging from 5.28 to 12.87), but the refined set showed consistently higher LDSC scores for all four phenotypes. For example, the P-value enrichment of the original set for bipolar disorder was 0.052, while the condensed enhancer set increased the statistical power by about 10x (P-value < 0.005). Even with only 12.6% of the coverage, the refined set improved the overall quality of the annotations and obtained a range of 2-to 10-fold increases in GWAS enrichment compared to the original set of annotations.

We believe the compactness of our refined annotation accounts for the increase in statistical power compared to the original set. Because of Grad-CAM, we are able to remove regions of variable lengths that do not contribute to the identity of an enhancer and shrink the prediction to 12.6% of its original size. Therefore, the refined annotations are more condensed and are more suitable for a wider range of analyses because many calculations, such as functional validation, require compact definitions to obtain statistical power.

## 4 Discussion

Here, we propose a *<u>De</u>*ep-learning framework for *<u>Co</u>*n*<u>de</u>*nsing enhancers and refining boundaries with large-scale functional assays (DECODE). Our model has two distinct parts: a binary classifier and an object detection module, both of which provide substantial benefits over previous methods.

For the binary classifier, we trained a deep learning model on direct readouts of human enhancer data to classify enhancer windows based on common epigenetic profiles. The classifier outperformed the *state-of-the-art* method in predicting cell-type-specific enhancers by using a larger set of training data and a deep learning model. CNNs have a larger capacity to learn non-linear, complex feature interactions compared to previous linear methods. We also emphasize that our deep learning-based DECODE model will benefit from the rapid development of novel functional characterization assays (e.g., MPRA and CRISPR-based screens) and the exponential growth of training data to further improve the accuracy and performance in enhancer predictions.

In addition to an improvement in prediction accuracy, our DECODE model also has a unique boundary detection module via Grad-CAM, which is not found in previous methods. The resultant feature-wise importance scores increase the interpretability by visualizing feature prioritization, while the position-wise importance scores can be used to condense the coarse enhancer annotations to the core functional regions. In particular, we show that our compact enhancer definitions have strong regulatory impact and are essential for disease causal variant mapping in disease studies.

In summary, we introduce a powerful tool that could be widely deployed for enhancer discovery. With corresponding epigenetic features, DECODE can not only accurately predict the existence of enhancers in any given genomic region, but also pinpoint the core functional regions, which greatly facilitates variant mapping and interpretation.

## Funding

This work was supported by the NIMH grant K01MH123896 and the NIH grant U01MH116492.

## Conflict of Interest

none declared.

## Appendix

**Table 1.** Data availability matrix from ENCODE. Data was available for all four histone marks across all cell types, but ATAC-seq was only available for K562 and HepG2 cells.

| Cell Type | STARR-seq | ATAC-seq | DNase-seq | H3K27ac ChIP-seq | H3K4me3 ChIP-seq | H3K4me1 ChIP-seq | H3K9ac ChIP-seq |
| --- | --- | --- | --- | --- | --- | --- | --- |
| K562 | ✓ | ✓ | ✓ | ✓ | ✓ | ✓ | ✓ |
| HepG2 | ✓ | ✓ | ✓ | ✓ | ✓ | ✓ | ✓ |
| A549 | ✓ |  | ✓ | ✓ | ✓ | ✓ | ✓ |
| HCT116 | ✓ |  | ✓ | ✓ | ✓ | ✓ | ✓ |
| MCF-7 | ✓ |  | ✓ | ✓ | ✓ | ✓ | ✓ |

## Notes

### Competing Interest Statement

The authors have declared no competing interest.

### Summary of Updates

Added code availability links

